# Genome-wide detection of positive and balancing selection signatures shared by four domesticated rainbow trout populations (*Oncorhynchus mykiss)*

**DOI:** 10.1101/2022.12.08.519621

**Authors:** K. Paul, G. Restoux, F. Phocas

**Affiliations:** Université Paris-Saclay, INRAE, AgroParisTech, GABI, 78350 Jouy-en-Josas, France

**Keywords:** Runs of Homozygosity, Extended Haplotype Homozygosity, domestication, fitness, selection, fish.

## Abstract

Evolutionary processes leave footprints across the genome over time. Highly homozygous regions may correspond to positive selection of favourable alleles, while maintenance of heterozygous regions may be due to balancing selection phenomena. We analyzed 176 genomes coming from 20 sequenced US fish and 156 fish from three different French lines that were genotyped using a HD Axiom Trout Genotyping 665K SNP Array. Using methods based on either Run of Homozygosity or Extended Haplotype Homozygosity, we detected selection signals in four domesticated rainbow trout populations. Nine genomic regions composed of 253 genes, mainly located on chromosome 2 but also on chromosomes 12, 15, 16, and 20, were identified under positive selection in all four populations. In addition, four heterozygous regions containing 29 genes putatively under balancing selection were also shared by the four populations and located on chromosomes 10, 13, and 19. Whatever the homozygous or heterozygous nature of the region, we always found some genes highly conserved among vertebrates due to their critical roles in cellular and nuclear organisation, embryonic development or immunity. We identify new promising candidate genes involved in rainbow trout fitness, as well as genes already detected under positive selection in other fishes (*auts2, atp1b3, zp4, znf135, igf-1α, brd2, col9a2, mrap2, pbx1, emilin-3*). These findings represent a genome-wide map of signatures of selection common over rainbow trout populations, which is the foundation to understand the processes in action and to identify what kind of diversity should be preserved, or conversely avoided in breeding programs, in order to maintain or improve essential biological functions in domesticated rainbow trout populations.

## 1 Introduction

Any population, whether animal or plant, wild or domesticated, evolved through continuous and cumulative changes over time (Wright, 1931). It relies on various evolutionary forces, mutation, migration, selection, and genetic drift, whose relative effects may vary depending on population history and structure. For example, genetic drift is more substantial when the effective population size is small and randomly induces fixation of alleles, which may lead to degeneration and extinction due to the fixation of deleterious alleles in small populations (Smith & Haigh, 1974). When modifications of environmental conditions occur, allele frequencies will change to a new relevant equilibrium, as a result of natural selection. Indeed, favorable alleles in a particular environment due to either new mutations or standing variation, will be positively selected. In wild populations, favourable alleles are generally affecting fitness through individual survival, mating, or fertility (East, 1918; Fisher, 1958). Natural selection can also act by negative (or purifying) selection that hinders the spread of deleterious alleles (Charlesworth et al., 1995). These two processes tend to reduce the genetic diversity at the target genes but had different effect on the genome, positive selection leading to stronger selection signatures (selective sweep) than negative one. Conversely, the population’s polymorphism can be actively maintained in some rare genomic regions through balancing selection that keeps an equilibrium in the frequencies of alleles. The two main biological causes of balancing selection are heterozygote advantage at a single locus, known as overdominance effect, and frequency-dependent selection with a rare-allele advantage that tends to restore a frequency equilibrium between alleles at the population level (Charlesworth, 2006, Fijarczyk & Babik, 2015).

Domestication is the evolutionary process of genetic adaptation over generations of a wild population to handling by humans and breeding in captive environments (Darwin, 1859, 1868; Price, 1984). During domestication, humans exert artificial selection pressure on the initial population by choosing and organizing the reproduction of the most adapted individuals to cohabitation or more globally to those whose aptitudes correspond the best to their expectations (Price, 1999; Russell, 2002), such as a less fearfulness of humans (Price, 2002; Harri et al., 2007). Domestication induces severe genetic bottlenecks due to the selection and reproduction of only a few adapted animals from the wild population. Thus, many genetic evolutionary processes, such as selection, genetic drift, and inbreeding, have a significant role in the evolution of farmed animal populations (Helmer, 1992; Mignon-Grasteau et al., 2005). The domestication process affects life history traits due to changes in morphological, physiological, reproductive, behavioural, and immune functions (Mignon-Grasteau et al., 2005; Pulcini et al., 2013; Milla et al., 2021 for review in fishes) compared to their wild relatives (Darwin, 1859, 1868). Wilkins et al. (2014) suggest that these specific modifications, called domestication syndrome, may be due to mild deficit of neural-crest cells during embryonic development in domesticated animals. In addition, both natural and artificial selection in domesticated species leaves footprints across the genome, known as selection signatures, which can point to regions harboring essential genes for domestication or natural fitness (Dobney & Larson, 2006; Qanbari & Simianer, 2014; Wright, 2015).

Compared to domestication in terrestrial animals (Mignon-Grasteau et al., 2005), fish domestication is recent and was first documented with carp about 2,000 years ago. The precise date and location (Neolithic China or at the Roman period in Central and East Europe) of the carp domestication are still debated (Balon, 1995; Balon, 2004). However, most farmed fish species have only been domesticated since the last century. The rainbow trout is native to the Pacific drainages of North America and to Kamchatka in Russia and its domestication started in the 1870s in California (Hershberger, 1992; Gall & Crandel, 1992). It was then introduced in Western Europe at the beginning of the 20th century (Fabrice, 2018).

Numerous studies have been carried out over the last ten years to detect signatures of selection in farmed fish species (Channel Catfish: Sun et al., 2014; Atlantic salmon: Mäkinen et al., 2015; Gutierrez et al., 2016; Liu et al., 2016; Pritchard et al., 2018; López et al., 2019; Carp: Su et al., 2018; Nile Tilapia: Hong et al., 2015; Cádiz et al. 2020; Yu et al., 2022; Rainbow trout: Cádiz et al., 2021; Coho salmon : López et al., 2021; Australasian snapper: Baesjou & Wellenreuther, 2021; Brown trout: Magris et al., 2022) in order to identify genomic regions involved in recent adaptation or domestication processes (Smith & Haigh, 1974; Pennings & Hermisson, 2006). In this study, we were interested in farmed rainbow trout populations as it is one of the oldest farmed fish and the analysis of genes under either positive or balancing subsequent selection in. Indeed, only a few studies on selection signatures were performed in rainbow trout. Three of them only focused on wild populations and showed signatures of selection linked to life-history variation, egg development, spawning time (Martínez et al., 2011), immune response (Limborg et al., 2012), and smoltification (Weinstein et al., 2019). The first study in domesticated rainbow trout was performed on a single Chilean population (Cádiz et al., 2021) genotyped with a 57K SNP array; identified signatures of selection were associated with early development, growth, reproduction and immune system. Recently, a high-density array (665K SNPs) was developed for rainbow trout (Bernard et al., 2022), allowing us to potentially more accurately detect signatures of selection and to compare them across various domesticated rainbow trout populations. The existence of signatures of selection shared by farmed populations from different geographical areas is essential to understand the importance of genetic diversity in several genomic regions in rainbow trout and then to identify genes having key roles in either the domestication process or fitness because conserved by all populations (Bruford et al., 2003; Yáñez et al., 2022).

Various approaches have been developed to reveal selection signatures within population based on site frequency spectrum, linkage disequilibrium (LD), or reduction of local variability (Vitti et al., 2013; Saravanan et al., 2020). Among these approaches, we will use two strategies, one based on the reduction of local variability using Run of Homozygosity (ROH) metrics and the second one relying on allele frequencies and the extent of LD based on Extended Haplotype Homozygosity (EHH). ROH is a large homozygous stretch in the genome of an individual inherited from a common ancestor to his parents (McQuillan et al., 2008; Purfield et al., 2012), while EHH measures the extent of shared haplotypes through the association between a single core haplotype and multiple loci at various distances from the core region (Sabeti et al., 2002). In our study, we considered four populations: one INRAE experimental line (with no intentional selection), two French selected lines from two different breeding companies, and a pooled American population gathering samples from one wild river and four hatchery populations, all coming from the North-West of the USA and closely genetically linked (Gao et al., 2018). Our work aimed to discover the main genomic regions sharing strong homozygosity (positive selection) or heterozygosity (balancing selection) across the four rainbow trout populations and to get further insights into the nature of genes spanning these regions.

## 2 Material and methods

### 2.1 Populations

Three French populations were considered: 14 breeding females from the INRAE synthetic line SY and, 90 and 72 females from two selected lines LB and LC from the breeding companies “Bretagne Truite” (Plouigneau, France) and “Viviers de Sarrance” (Sarrance, France) respectively. The SY was developed by intercrossing several domesticated lines of rainbow trout, in order to create a population with a large diversity (D’Ambrosio et al., 2019).

In addition, we considered an American pooled population, hereafter named HA, using the whole genome sequence data of 20 fishes obtained by Gao et al. (2018). The sampling strategy consisted in collecting DNA from 4 individuals in each of five locations from the North-West of the USA: wild fish from Elwha River, and farmed fish from Dworshak, L. Quinault, Quinault, and Shamania hatcheries. We pooled together the 20 individuals, as these five populations were genetically close to each other (Supplementary Figure 1; Gao et al., 2018) and greatly distant from the three French populations (Figure 1).

**FIGURE 1.**
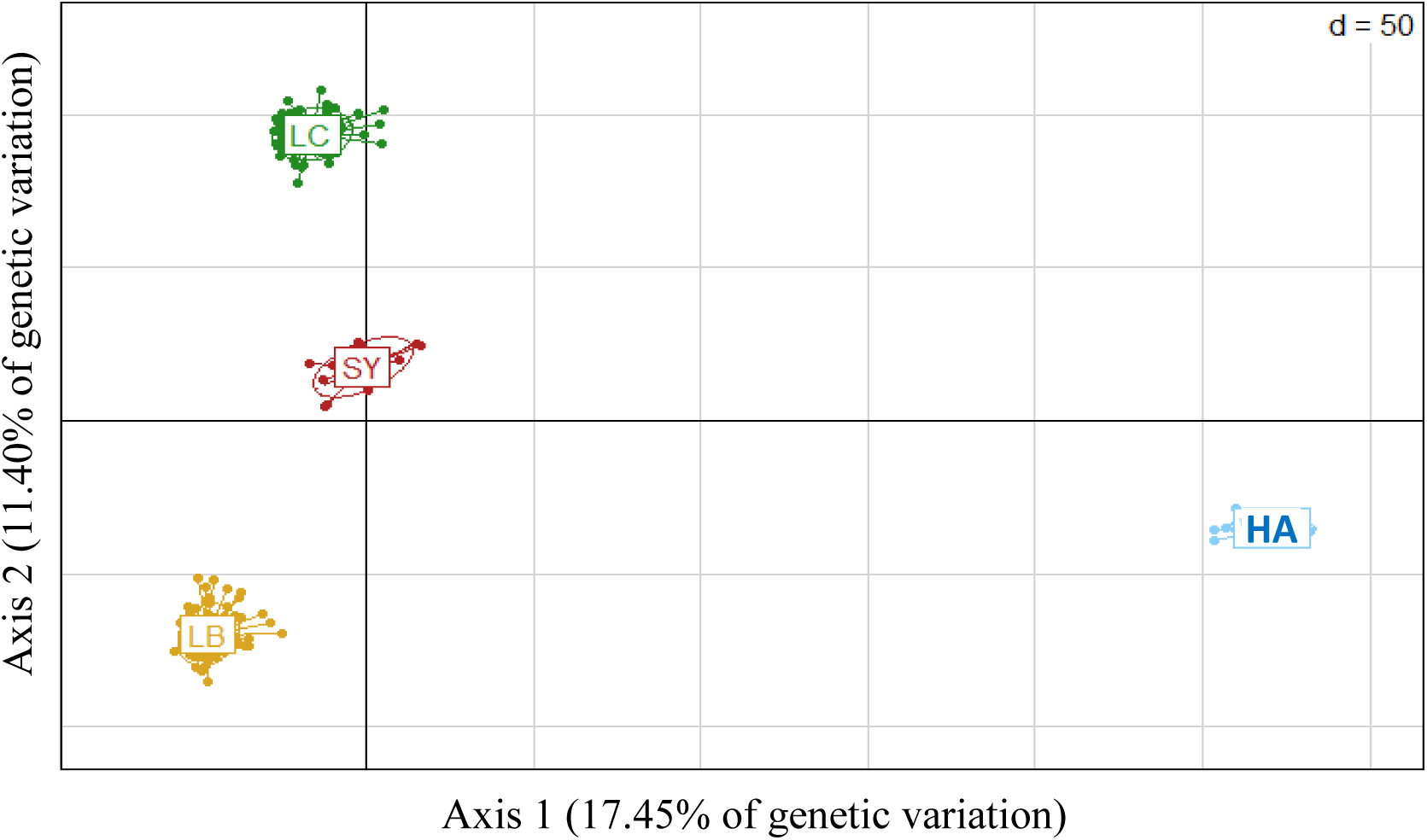
Principal component analysis (PCA) of the genetic diversity of the four rainbow trout populations (LB, LC, SY, and HA) based on 546,903 SNPs.

### 2.2 Genotyping and quality control

High-density genotypes were obtained at the INRAE genotyping Platform Gentyane (Clermont-Ferrand, France) for all the 176 French samples using the Affymetrix 665K SNP array recently developed for rainbow trout (Bernard et al., 2022). We only considered the genotypes for the 576,118 SNPs of the Rainbow Trout Axiom® 665K SNP array that were positioned on the Arlee genome (GCA_013265735.3, Gao et al., 2021; Bernard et al., 2022). From the whole-genome sequence information of the 20 American samples (Gao et al., 2018), we extracted the genotypes for the same 576,118 SNPs of the HD chip.

Among the 177 French genotyped fish, 19 individuals with more than 30% identity-by-state (IBS) with other individuals were removed from the dataset. We thus kept for the analysis 76, 67, 20, and 14 fish sampled from LB, LC, HA, and SY populations, respectively.

Then, SNP quality control was performed using PLINK v1.9 software (Chang et al., 2015). Note that, to avoid limitations due to the low number of individuals in SY, quality filters were made considering LC and SY together, as both populations were genotyped on the same SNP plate and are close genetically (D’Ambrosio et al., 2019). About 4,000 SNPs randomly distributed over all the genome were removed for all populations due to extreme deviation from Hardy-Weinberg equilibrium (p-value < 10-7). It allowed us to discared SNPs with high risk of wrong genotyping, in addition to the edit for SNP call rate lower than 97%. We retained 571,319 SNPs, 569,030 SNPs, and 573,793 SNPs on LB, ’LC-SY’, and HA populations, respectively. Finally, crossing the three SNP lists, we kept the 546,903 common SNPs for the analysis.

### 2.3 Genetic structure of the populations

Genetic differentiation between populations was measured with a pairwise Fst estimate using the VCFtools v0.1.13 software (Danecek et al., 2011). In addition, a principal component analysis (PCA) was performed with the R package *Adegenet* (function *glPca*) (Jombart & Ahmed, 2011) to visualize the genetic structure of the populations.

### 2.4 Runs of homozygosity

Runs of homozygosity (ROH) were identified for each fish using the PLINK v1.9 *homozyg* function (Chang et al., 2015) with the following options *‘--homozyg-kb 500 –homozyg-window-snp 40 –homozyg-snp 40 –homozyg-gap 500 –homozyg-density 40 –homozyg-het 1*’. ROH was defined by a sliding window with a minimum length of 500 kb containing at least 40 homozygous SNPs. This minimum number of homozygous SNPs was chosen using the formula described by Purfield et al. (2012) in order to limit the number of ROHs that might only occur by chance.

#### 2.4.1 Estimation of inbreeding coefficients

The individual inbreeding coefficients (F_ROH_) were calculated according McQuillian et al (2008) as

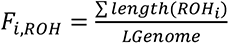

With ∑*length*(*ROH_i_*) the sum of ROH length in an individual i and *LGenome* the total length of the autosomal genome covered by SNPs.

#### 2.4.2 Identification of ROH islands

For each SNP, the number of individuals with this SNP included in a ROH was calculated in order to identify the regions of the genome that were frequently homozygous in each population, i.e. constituting ROH islands (Nothnagel et al., 2010). These ROH hotspots may then be considered as signatures of positive selection (Saravanan et al., 2021).

To allow the comparison of ROH islands across populations, we implemented population-specific thresholds based on the ROH occurrence to define ROH islands, as proposed in many studies (Purfield et al., 2017; Mastrangelo et al., 2017; Zhang et al., 2018; Peripolli et al., 2018; Grilz-Seger et al., 2018; Gorssen et al., 2021; Illa et al., 2022). The number of individuals corresponding to the top 5% of SNPs most often found in a ROH within each population was chosen as a threshold to define a ROH island.

These top 5% values were equivalent to 35, 27, 5, and 10 individuals for LB, LC, SY, and HA, respectively. Values chosen within each population corresponded to 48.6%, 40.3%, 35.7%, and 50% of individuals with a ROH in LB, LC, SY, and HA, respectively. In addition, two close SNPs in the top 5% were considered in the same ROH island if there were separated by a distance lower than 500 kb with less than 40 SNPs in the gap stretch. The ROH island was delimited by a number of individuals, with ROH falling below the top 10% of the SNPs, which correspond to 30, 22, 3, and 7 individuals for LB, LC, SY, and HA populations, respectively.

#### 2.4.3 Detection of balancing selection signals based on regions without ROH

We used the ROH occurrence information per SNP to detect extreme heterozygous regions, i.e. without ROH. In these regions, we have an enrichment of heterozygous SNP relative to the genome-wide prevalence that may be due to balancing selection phenomena (Szpiech et al., 2013).

Applying the same criteria as for defining ROH, the minimal size and number of SNPs to define a heterozygous region were fixed to 500 kb and 40 SNPs, respectively. Moreover, two successive SNPs were considered in the same region if they were separated by a distance lower than 50 kb. A region was detected in extreme heterozygosity if less than 5% of individuals (per population) have SNPs in ROH in the region, corresponding to a maximum of respectively 4 and 3 individuals with a ROH in LB and LC populations, and to 0 individual with a ROH in SY and HA.

### 2.5 Detection of signatures of selection based on Extended Haplotype Homozygosity (EHH)

For a given core allele, the EHH is defined as ‘the probability that two randomly chosen chromosomes carrying the core haplotype of interest are identical by descent for the entire interval from the core region to the point x’ (Sabeti et al., 2002). EHH measures the association between a single allele from the study locus (the core region) with multiple loci at various distances x (Sabeti et al., 2002). The iHS (Integrated Haplotype Homozygosity Score) proposed by Voight et al. (2006) aims to compare the integrated EHH profiles obtained for a SNP in the ancestral versus derived states. An extreme value of iHS corresponds to a positive selection because a core haplotype with unusually high EHH and high frequency in the population indicates the presence of a mutation that has spread through the population faster than the haplotype broke down.

EHH methodology requires haplotype information. Thus, genotype data must be phased before their calculation. We used FImpute3 (Sargolzaei et al., 2014) to phase the genotypes of the study females, considering all parents (including our study females) and offspring genotyped in LB, LC, and SY populations for different purposes (see respectively Prchal et al., 2022, Lagarde et al., 2022 and Paul et al., 2022). All parents (except 8 SY sires) were genotyped with the HD chip (Bernard et al., 2022), while offsprings (and 8 SY sires) were genotyped with a 57K chip only (Palti et al., 2015). Information used for phasing is given in Table 1. Due to the lack of genotyped offsprings, only the HD genotypes information was used to phase the genotypes of the HA population.

**TABLE 1.** Data information used to phase the HD genotypes of the study females that belong to the parental cohorts. Number of individuals and SNPs available after quality control

| Line | Status of individuals | Number of individuals | Number of SNP used |
| --- | --- | --- | --- |
| LB | parents | 288 | 571,319 |
|  | offsprings | 1,297 | 29,091 |
| LC | parents | 173 | 569,03 |
|  | offsprings | 1,350 | 30,379 |
| SY | parents (dams + 1 sire) | 16 | 569,03 |
|  | offsprings (+ 8 sires) | 866 | 32,725 |

Once phasing was performed, the *rehh* R package (Gautier & Vitalis, 2012; Gautier et al., 2017) was used to conduct EHH-based analyses. EHH detection was stopped when the EHH value declined under 0.1 or when the gap between two consecutive SNPs was higher than 20 kb (*scan_hh* function with the following options: limehh = 0.1; maxgap=20 kb).

#### 2.5.1 Cross population Extended Haplotype Homozygosity (XP-EHH)

From EHH information, we used the XP-EHH statistic (*ies2xpehh* function) to compared the integrated EHH profiles (iES), two by two, between a French (popA) and the HA (popB) populations at the same focal SNP (Sabeti et al., 2007) as:

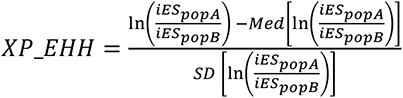

The median (*Med*) and standard deviation (*SD*) of ln(iES_A_/iES_B_) were computed over all the analysed SNPs.

#### 2.5.2 Integrated Haplotype Homozygosity Score (iHS)

In the same way, we used the iHS test (Voight et al., 2006) to evaluate the evidence of positive selection based on haplotype frequencies in a single population, using the *ihh2ihs* function of the package *rehh*. This statistic was based on the log-ratio of the integrated EHH (iHH) for haplotypes with the ancestral (A) *versus* the derived (D) alleles and was computed for each autosomal SNP as

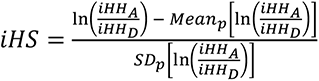

The average (*Mean_p_*) and standard deviation (*SD_p_*) of ln(iHH_A_/iHH_D_) were computed over all the SNPs with a derived allele frequency *p* similar to that of the core SNP. In our study, the ancestral allele state is unknown. Therefore, we assumed that the most frequent allele represents the ancestral state as proposed by Bahbahani et al. (2015).

#### 2.5.3 Detection of candidate regions

To detect candidate regions for signatures of selection based on the iHS test, we used *the calc_candidate_region* function of the R package *rehh* (Gautier & Vitalis, 2012). We considered windows of 500 kb across the genome containing at least 30 SNPs, with 10 kb of overlapping. A region was considered as under positive selection if at least one SNP had a log(p-value) > 4 and extreme iHS value *i.e.* iHS ≥ 2.5.

### 2.6 Identification of common regions under positive selection

ROH islands and regions identified by iHS were pooled within each population. Then, the intersection set of the regions identified by one or another method across the four studied populations was established. We eliminated an intersection from the study if one population does not have at least one SNP with an iHS≥ 2.5 or enough individuals with an ROH in the intersected region. So, only intersections containing either ROH island or extreme iHS (iHS ≥ 2.5) for the four populations were thus further analyzed.

### 2.7 Gene analysis

The genes annotated in the regions under positive or balacing selection were identified from the NCBI *Oncorhynchus mykiss* genome assembly (GCA_013265735.3). Gene symbols were checked, and, if necessary, familiar names were added using the information available from GeneCards (https://www.genecards.org/).

Gene ontology (GO) terms study was performed with ’g:profiler’ (Raudvere et al., 2019; https://biit.cs.ut.ee/gprofiler/gost) for the list of genes identified in the regions of interest. Percent identity of rainbow trout proteins with nine other vertebrate species (human, mouse, cow, goat, pig, chicken, zebrafish, medaka, and Atlantic salmon) was established using the blastp tool (Basic Local Alignment Search Tool on proteins).

## 3 Results

### 3.1 Genetic diversity within and across populations

The ROH statistics and inbreeding coefficients are presented in **Table 2** for all the populations. The average number of ROH per individual varied between 141 (SY) and 168 (LB). French selected lines had larger average sizes of ROH than populations SY and HA. The average inbreeding coefficients of HA individuals were between three (compared to SY) and five (compared to LB) times lower than those of the French lines.

**TABLE 2.** ROH statistics and inbreeding coefficients of the four studied populations (Standard deviations are indicated in brackets).

| Population | Average number of ROH | Average size of ROH (in kb) | Average $F_{ROH}$ |
| --- | --- | --- | --- |
| LB | 168 (14.6) | 2,770 (270.8) | 0.20 (0.02) |
| LC | 157 (15.9) | 2,485 (326.8) | 0.17 (0.03) |
| SY | 141 (33.5) | 1,860 (291.2) | 0.12 (0.05) |
| HA | 167 (65.6) | 1,433 (145.6) | 0.04 (0.03) |

Based on genome-wide Fst values, large differentiation of around 0.28 was observed between HA and any of the French populations (Table 3). In the PCA figure (Figure 1), the three French lines differed strongly from the American pooled populations, and the first two PCA axes explained 29% of the total genetic variation. In addition, Fst values indicated that all the French lines were moderately differentiated (0.104 – 0.122).

**TABLE 3.** Genome-wide Fst statistics derived two-by-two between the four populations.

|  | LC | LB | HA |
| --- | --- | --- | --- |
| SY | 0.104 | 0.122 | 0.275 |
| LC |  | 0.122 | 0.275 |
| LB |  |  | 0.289 |

Using the XP-EHH statistic, we identified 93, 105 and, 135 regions that strongly discriminated HA from LB, LC and SY, respectively. Among these regions, 34 regions were shared, spanning about 32 Mb in total over 21 chromosomes, and differentiated any of the French lines from the American HA pooled population (Supplementary information S1).

The distribution of the proportion of individuals having a ROH at each SNP position is presented in Figure 2. In average, ROH were more shared between individuals for selected lines (LB and LC, on average, 23.39% and 19.82% of individuals respectively) than for other populations (SY and HA, on average, 13.67% and 8.91% of individuals respectively). Probably linked to the composite nature of the HA population (5 sub-groups of 4 individuals), HA contained the lowest number of shared ROH among the individuals but also showed the most shared ROH among individuals.

**FIGURE 2.**
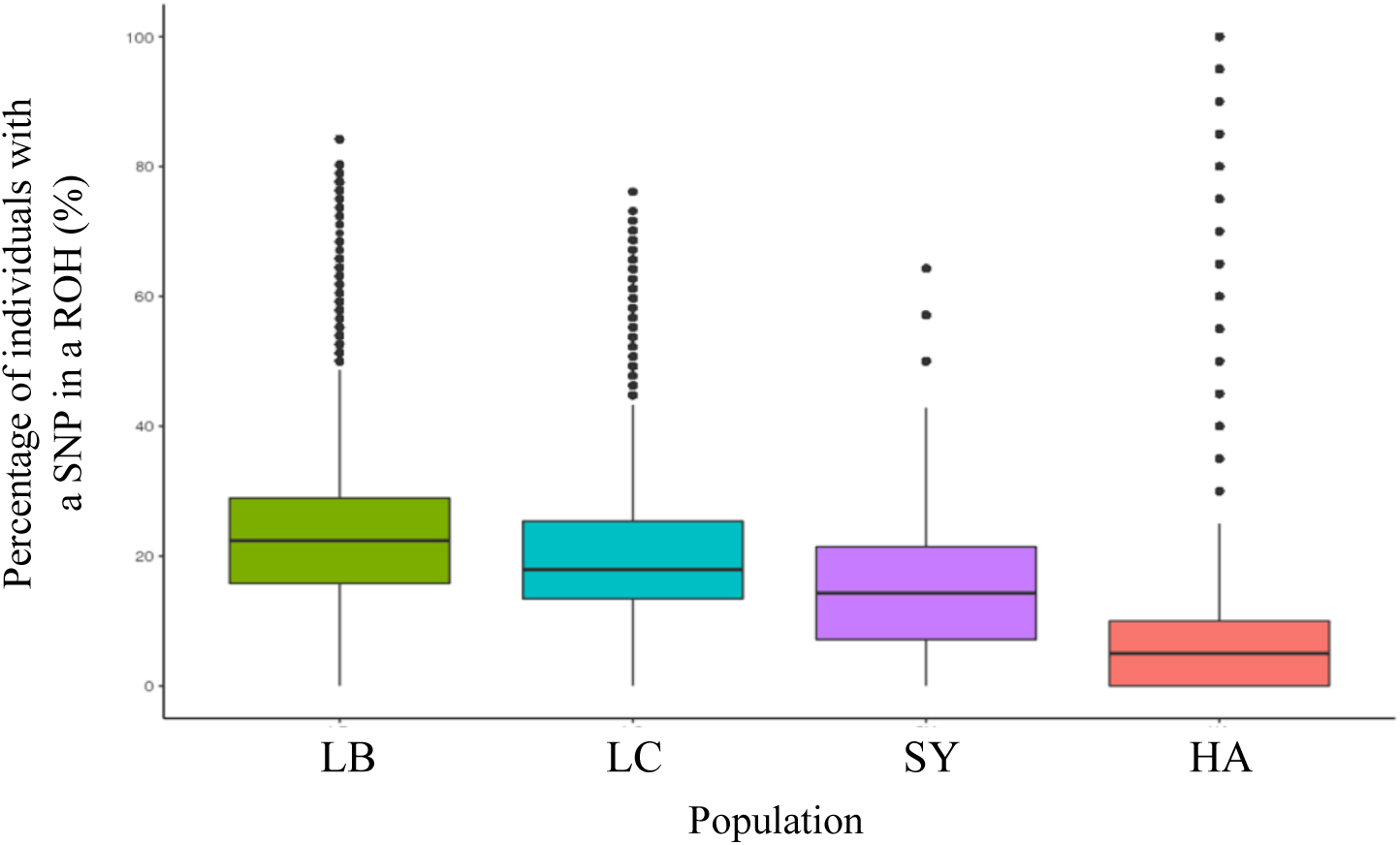
Box-plots of the occurrence of ROH (number of individuals having this ROH) per SNP for each rainbow trout population LB, LC, SY, and HA.

### 3.2 Signatures of positive selection

#### 3.2.1 ROH islands

The sharing of ROH among individuals, regardless of the population considered, was presented Figure 3. Eight ROH islands were shared by at least 2 populations, and a minimum of 50% of individuals concerned. However, only three of these regions were defined as ROH island in each of the 4 populations.

**FIGURE 3.**
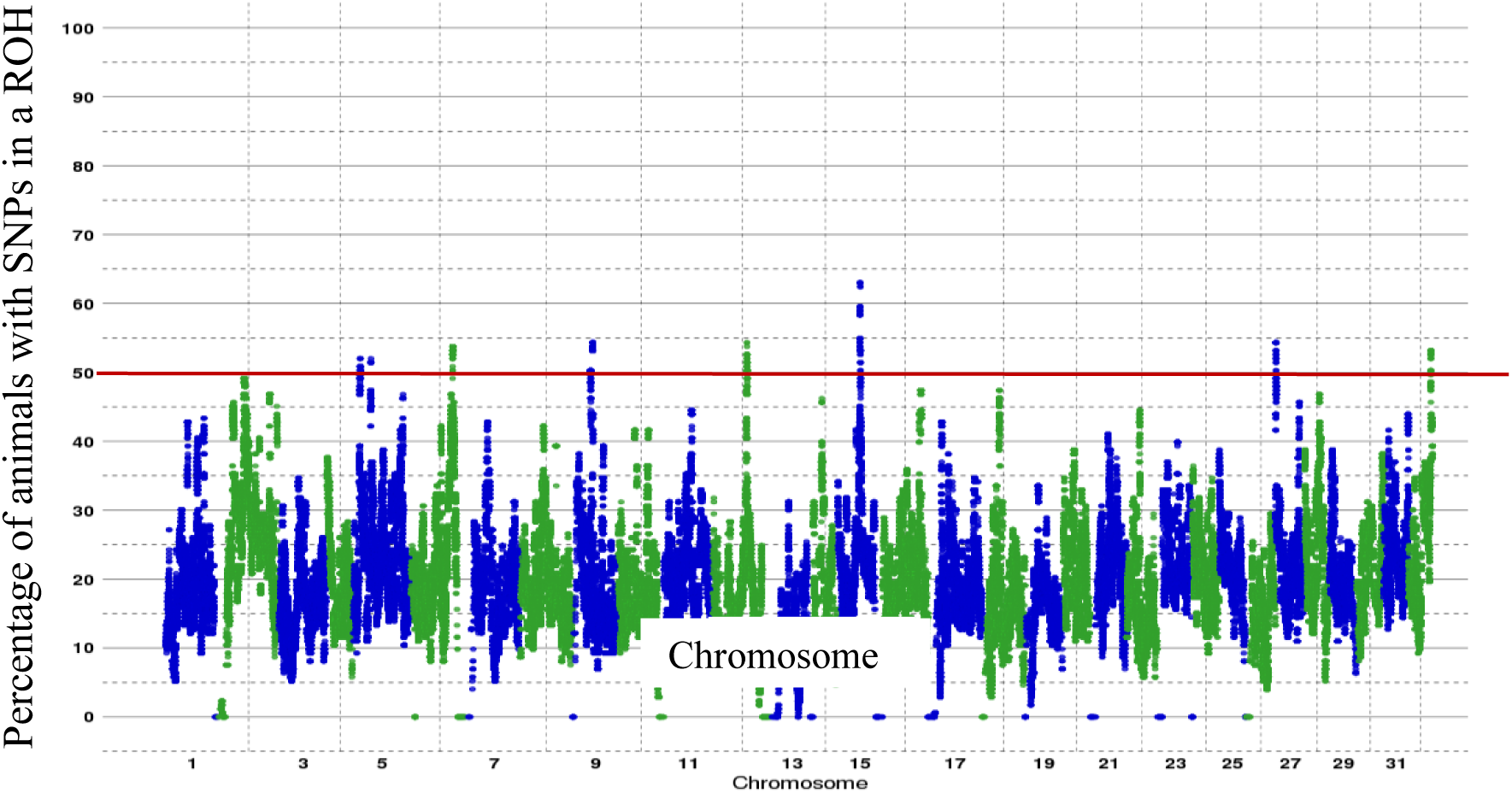
Manhattan plot of the occurrence of ROH per SNP across chromosomes (gathering all rainbow trout populations). The red line highlights the ROH islands.

We listed all ROH islands within each population which resulted in the identification of 270 ROH islands distributed among the four populations (Supplementary Informations S2 to S5, for LB, LC, SY, and HA, respectively). The ROH islands were not evenly distributed across populations and chromosomes. The average ROH island size was 2,737 kb, varying from 1,593 kb to 4,465 kb, depending on the population. The longest ROH island was observed in SY (21.4 Mb), while the shortest one was observed in LC (16.1 kb).

#### 3.2.2 Identifying selection signatures using iHS

The log(p-values) of iHS calculated along the genome are presented in Figure 4 for each population (all regions identified with *calc-candiate_region* are described in Supplementary Informations S6-S9). While numerous regions have been detected as under positive selection overall, fewer candidate regions were detected for the French lines (LB, LC and, SY) than for the Amercian pooled population (HA). The genome-wide highest estimated values of iHS were 8.97, 7.24, 5.67, and 9.09 for LB, LC, SY, and HA, respectively (with log(p-values) > 7.8).

**FIGURE 4.**
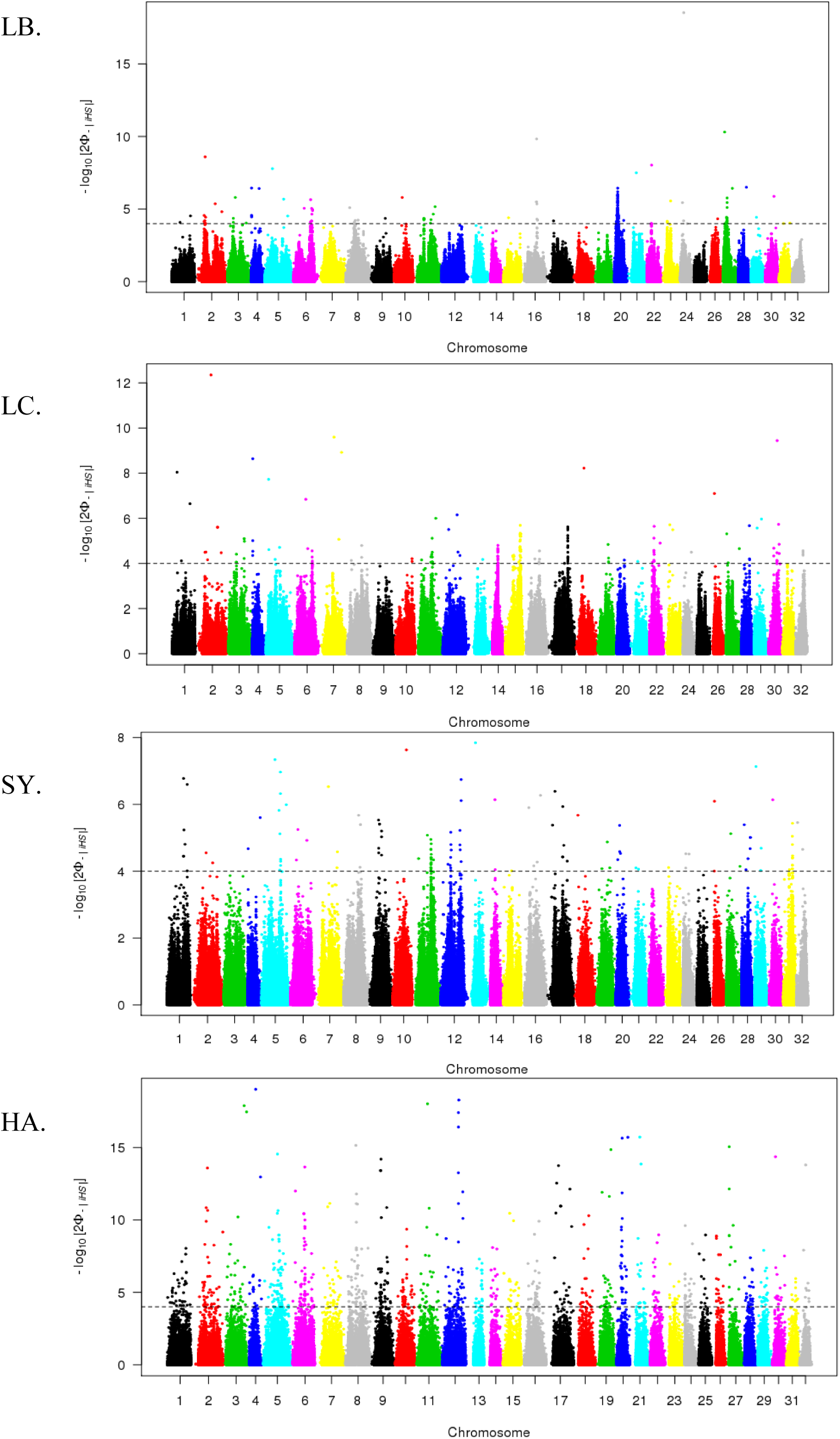
Genome-wide distribution of log(p-value) for standardized iHS for each population (LB, LC, SY, HA). The dashed line indicates the log(p-value) significance threshold set to 4 to identify regions under positive selection

In total, 72, 68, 76, and 54 ROH islands were identified in LB, LC, SY, and HA populations respectively (Figure 5). Using iHS statistics, 55, 69, 73, and 362 signatures of selection were detected for LB, LC, SY, and HA populations, respectively. Only 10.4%, 8.7%, 8.0%, and 5.6% of the regions were detected by both methods (ROH + iHS) for LB, LC, SY, and HA populations, respectively.

**FIGURE 5.**
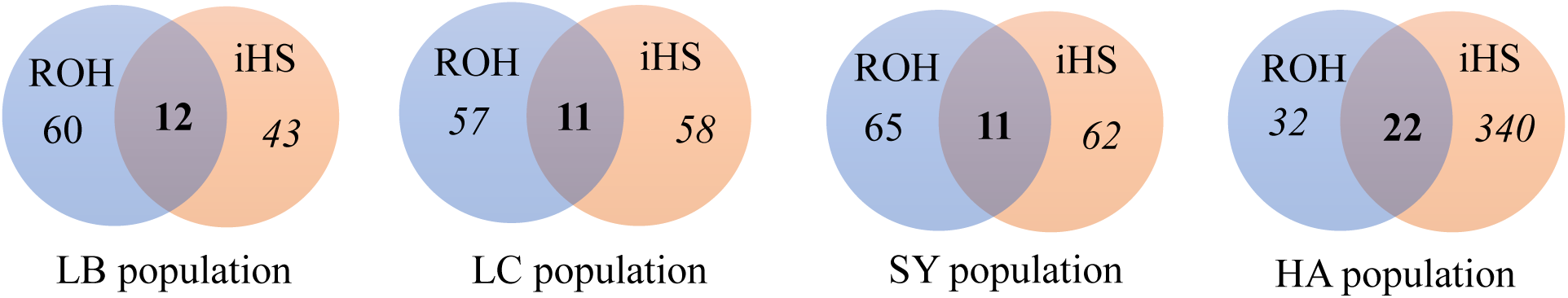
Venn Diagram of the number of regions identified as ROH island or iHS signature of selection for each rainbow trout population.

#### 3.2.3 Regions under positive selection shared by the four studied populations

Among the numerous regions identified for each population by either ROH or iHS methods, only nine regions were shared by the four studied populations (Table 4). The average size of these shared regions was 1135 kb. Five of them were located on chromosome 2, and the four other regions were on chromosomes 12, 15, 16, and 20, respectively.

**TABLE 4.** Homozygous regions under positive selection in the four studied populations.

| Region | CHR | Start (Mb) | End (Mb) | Size (kb) |
| --- | --- | --- | --- | --- |
| chr2_a | 2 | 25.40 | 26.30 | 900 |
| chr2_b | 2 | 31.60 | 34.20 | 2600 |
| chr2_c | 2 | 46.00 | 46.66 | 664 |
| chr2_d | 2 | 69.70 | 71.20 | 1500 |
| chr2_e | 2 | 88.46 | 89.34 | 878 |
| chr12_a | 12 | 57.97 | 59.10 | 1138 |
| chr15_a | 15 | 38.96 | 39.57 | 610 |
| chr16_a | 16 | 45.80 | 47.00 | 1200 |
| chr20_a | 20 | 19.10 | 19.83 | 726 |

Depending on the population, six regions were identified by either ROH or iHS metrics. Two regions, chr2_c and chr15_a, were only detected by ROH in all four populations, while a single region, chr16_a, was only identified through significant iHS statistics in all the four populations (Supplementary Information S10). The list of genes annotated in the nine shared genomic regions is given in Supplementary Informations S11.

### 3.3 Signatures of balancing selection

#### 3.3.1. Regions under balancing selection detected within population

In total, 14, 24, 158, and 265 hot spots of polymorphism (i.e. without ROH) were identified in LB, LC, SY, and HA populations, respectively. The numbers of heterozygous regions detected for SY and HA populations were drastically larger than those observed for the LB and LC selected lines. The average size of the detected heterozygous regions was 1,400 kb, varying from 1,086 kb to 1,828 kb, depending on the population.

Tables listing all heterozygous regions within each population are presented in Supplementary Informations S12 to S15, for LB, LC, SY, and HA, respectively.

#### 3.3.2. Regions under balancing selection shared by the four studied populations

A substantial lack of ROH was observed in four regions of all studied rainbow trout populations (Table 5). Two of them, chr10_a and chr19_a, were particularly small (53 kb and 70 kb, respectively), but still contained at least 20 SNPs. The region chr10_a only encoded one of the introns of the *ctnna2* (= catenin alpha 2) gene while chr19_a was composed of two genes, *smarca5* (=SWI/SNF-related matrix-associated actin-dependent regulator of chromatin subfamily A member 5) and *frem2* (= FRAS1-related extracellular matrix protein 2). A second heterozygous region on chromosome 19 was larger (163 kb) but contained a single annotated gene, *pou4f2* (= POU domain, class 4, transcription factor 2-like). A last region chr13_a spanned over 1,100 kb) on chromosome 13 and was composed of 25 genes. The list of genes annotated in the four shared genomic regions is given in Supplementary Informations S16.

**TABLE 5.** Highly heterozygous regions shared by the four studied populations.

| Region | CHR | Start (Mb) | End (Mb) | Size (kb) | SNP number | SNP density per Mb |
| --- | --- | --- | --- | --- | --- | --- |
| chr10_a | 10 | 56.314 | 56.366 | 53 | 20 | 379 |
| chr13_a | 13 | 46.959 | 48.071 | 1,112 | 446 | 401 |
| chr19_a | 19 | 10.753 | 10.823 | 70 | 24 | 342 |
| chr19_b | 19 | 11.354 | 11.517 | 163 | 52 | 319 |

### 3.4 Identification and roles of genes underlying the regions under selection across all populations

#### 3.4.1. Common homozygous regions under positive selection

The nine common homozygous regions contained a total of 253 genes (listed in Supplementary Information S11). A gene ontology (GO) study was performed and showed a significant over-representation (p-value < 0.01) among the 253 genes of functions related to the following GO terms: membrane (GO:0016020, CC: Cellular Component*, p-value = 1.3e10^-5^*), intrinsic and integral component of membrane (GO:0031224; GO:0016021, CC, *p-value = 0.001/0.005*), ion binding (GO:0043167, MF: Molecular Function*, p-value = 0.002*), and nuclear speck (GO:0016607, CC*, p-value = 0.008*).

Among the nine studied regions, the three regions chr2_a, chr2_c, and chr15_a, that contain less than ten genes annotated in each, were analyzed in further detail to accurately define the roles of underlying genes. The 17 genes located in these three regions are listed in Table 6 with their associated biological functions. These genes play key roles in protein transduction/maturation, genome stability, embryonic development, growth, energetic function, reproduction, or immune function. In addition to this list of genes, a subset of 15 genes in the six other homozygous regions already identified as signatures of selection in the literature were further studied in terms of their biological functions. Detailed information for these genes is also given in Table 6.

**TABLE 6.**
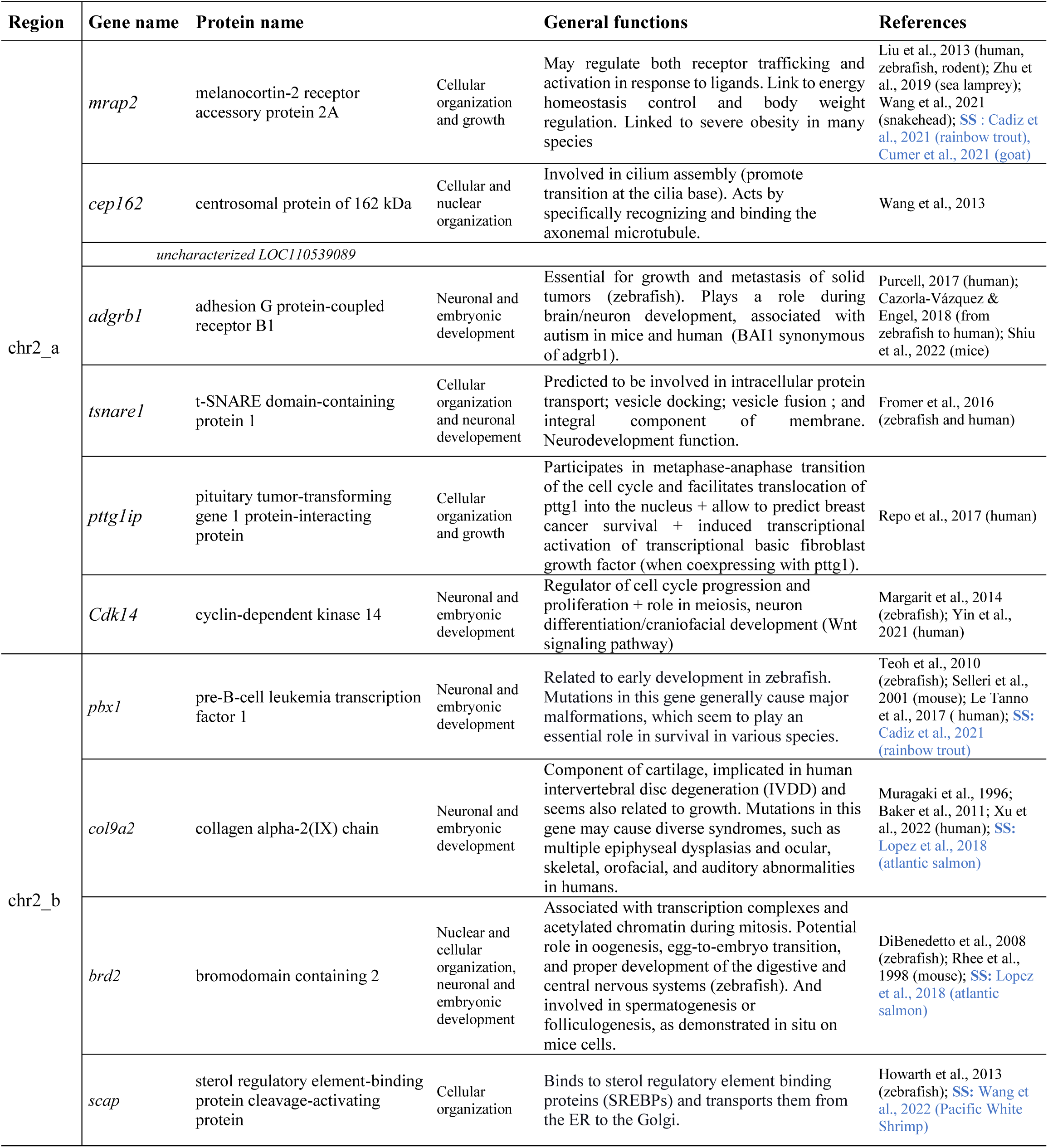
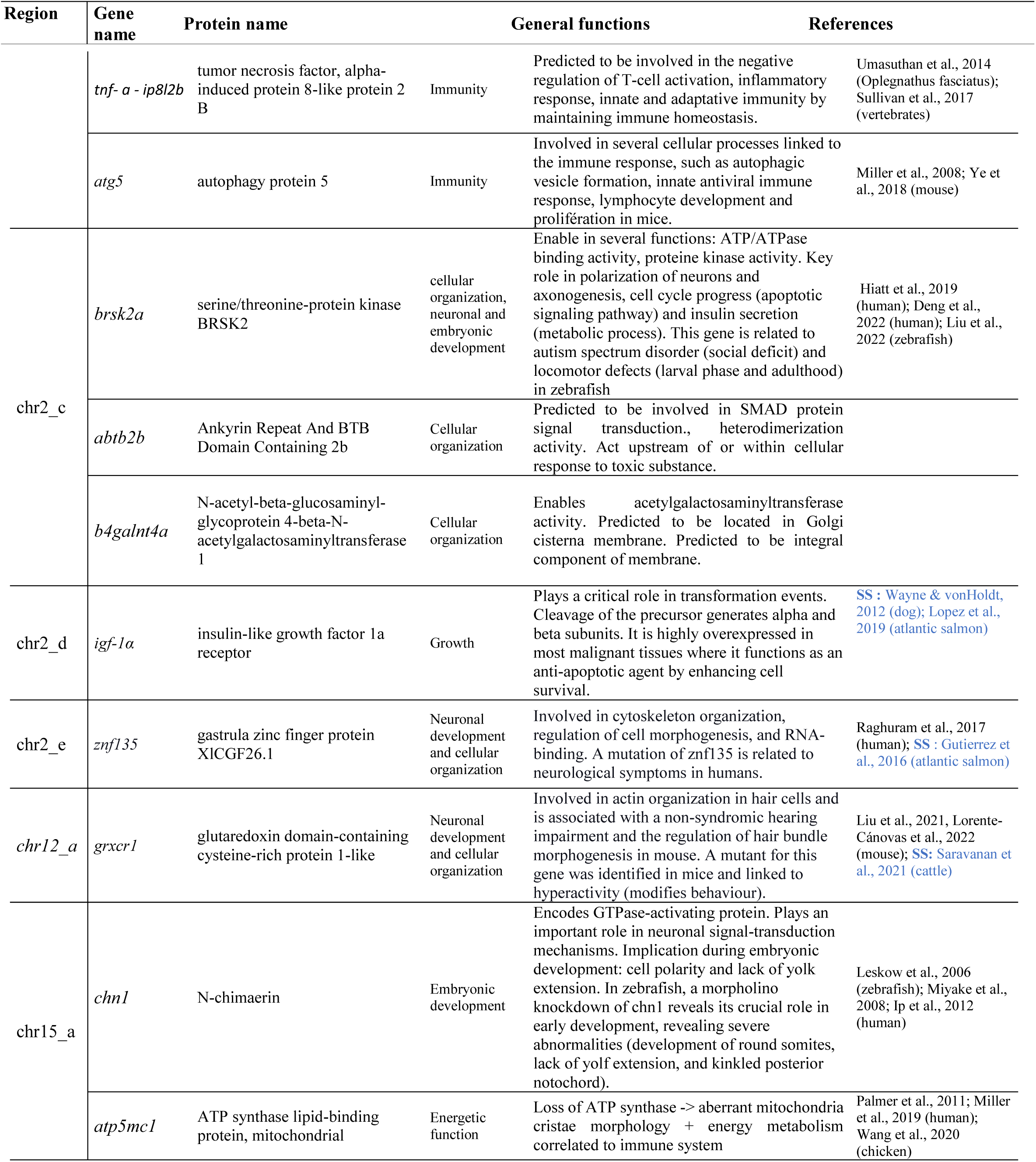
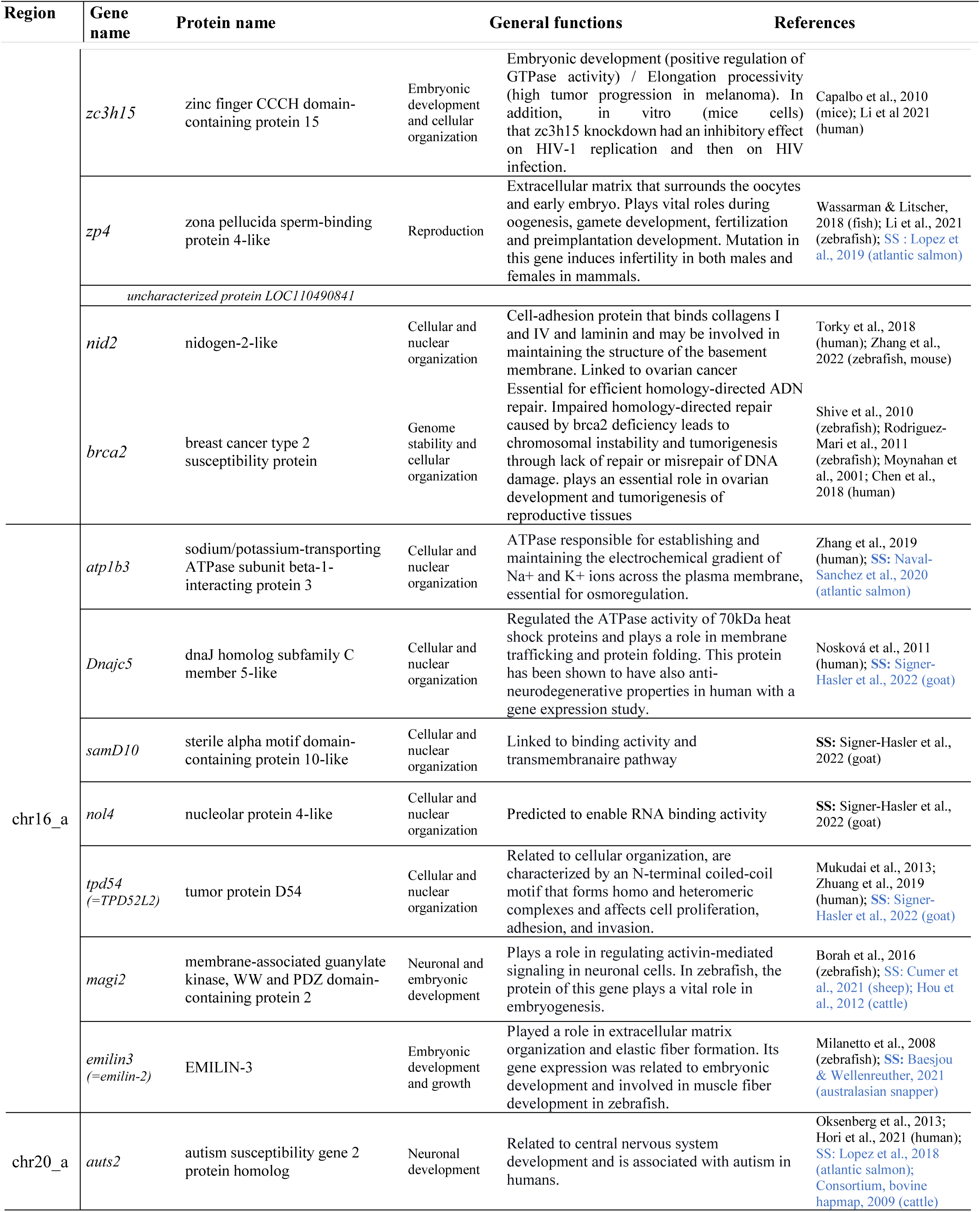
List and functions of the 17 genes annotated in three homozygous regions (chr2_a, chr2_c and chr15_a) shared by the four rainbow trout populations, and the 15 genes in the six other regions already identified as signatures of selection in the literature. *SS : Identify by signature of selection in that study

We studied the degree of protein identity among 10 vertebrate species for all the 17 genes of regions chr2_a, chr2_c, and chr15_a (Table 7), considering a protein as highly conserved if its identity between rainbow trout and other species was higher than 85%. Except for the proteins linked to *cep162* (centrosomal protein of 162 kDa) and *zp4* (zona pellucida sperm-binding protein 4-like) genes, all other proteins were highly conserved at least between the two studied salmonids. In each of the three regions, one or two genes were highly conserved across the all ten study species: in chr2_a, rainbow trout *cdk14* (cyclin-dependent kinase 14) protein had a percent identity between 86 and 99.6% with the other species; in chr2_c, rainbow trout *brsk2a (*serine/threonine-protein kinase *brsk2*) protein had between 92 and 96.3 % of percent identity with the other species; in chr15_a, two genes, *chn1*(n-chimaerin) and *atp5mc1* (ATP synthase lipid-binding protein, mitochondrial), also had protein percent identity ranging from 85% to 98% depending of the species.

**TABLE 7.**
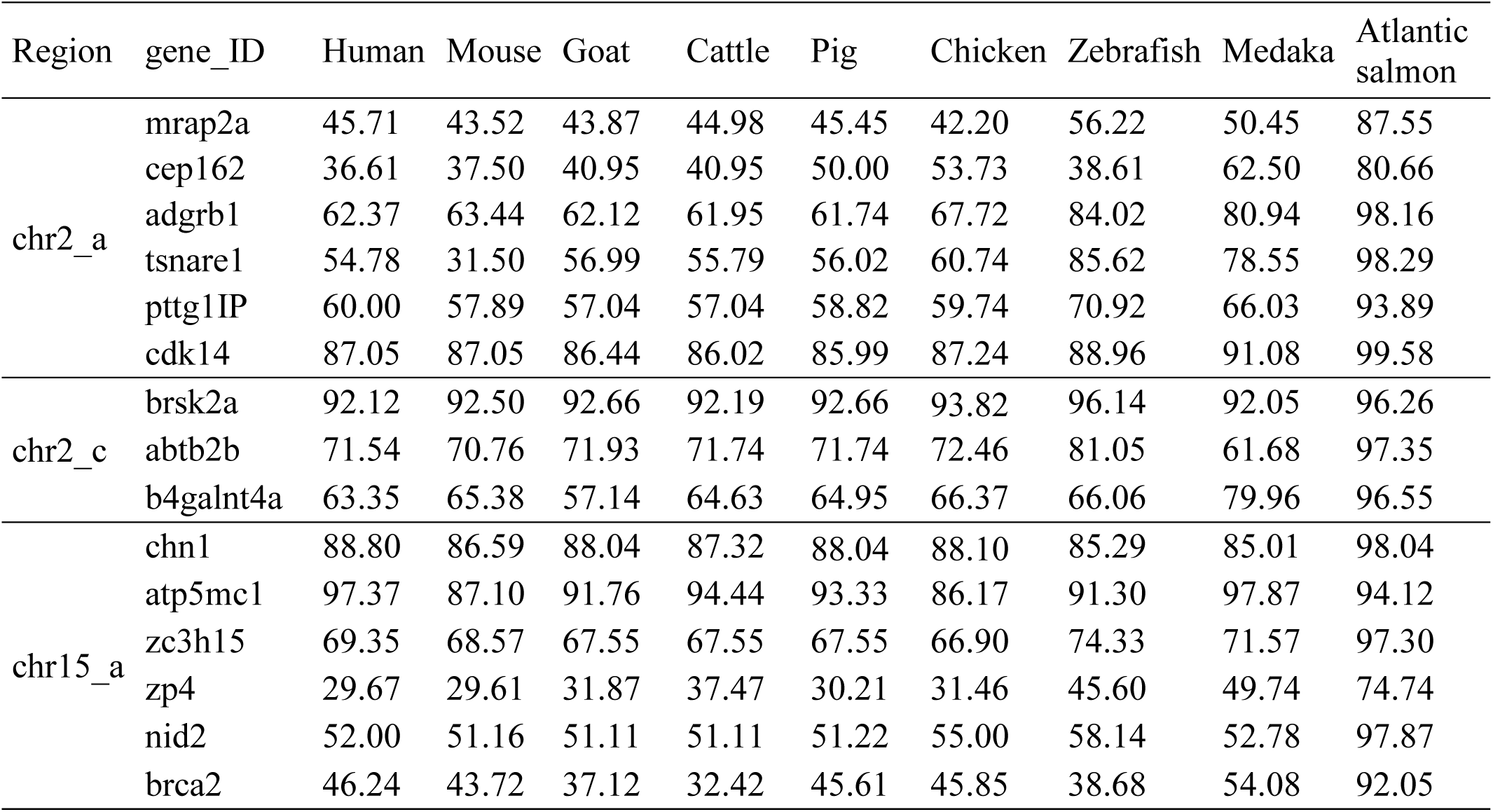
Percentage of protein identity between rainbow trout and nine other vertebrate species for all genes annotated in homozygous regions chr2_a, chr2_c and chr15_a.

Some other rainbow trout proteins (*tsnare1,* t-SNARE domain-containing protein 1*; pttg1IP,* pituitary tumor-transforming gene 1 protein-interacting protein) were conserved to a lesser extent (minimum 65% of percent identity) with the three other fish species, some being also conserved at least with chicken (*adgrb1,* adhesion G protein-coupled receptor B1*;b4galnt4a,* N-acetyl-beta-glucosaminyl-glycoprotein 4-beta-N-acetylgalactosaminyltransferase 1) or even with all the nine study species (*zc3h15,* zinc finger CCCH domain-containing protein 15).

#### 3.4.2. Common heterozygous regions under balancing selection

The four common heterozygous regions (Table 5) contained 29 genes (listed in Supplementary Information S16). A gene ontology (GO) terms study showed no significant over-representation of specific GO terms.

The degree of protein percent identity among 10 various vertebrate species for these 29 genes are presented in supplementary information S17.

Regions chr10_a, chr19_a, and chr19_b contained only a few genes and were then analyzed in further detail to accurately determine the role of underlying genes (Table 8). These genes play key roles in cellular and nuclear organisation and in embryonic development.

**TABLE 8.**
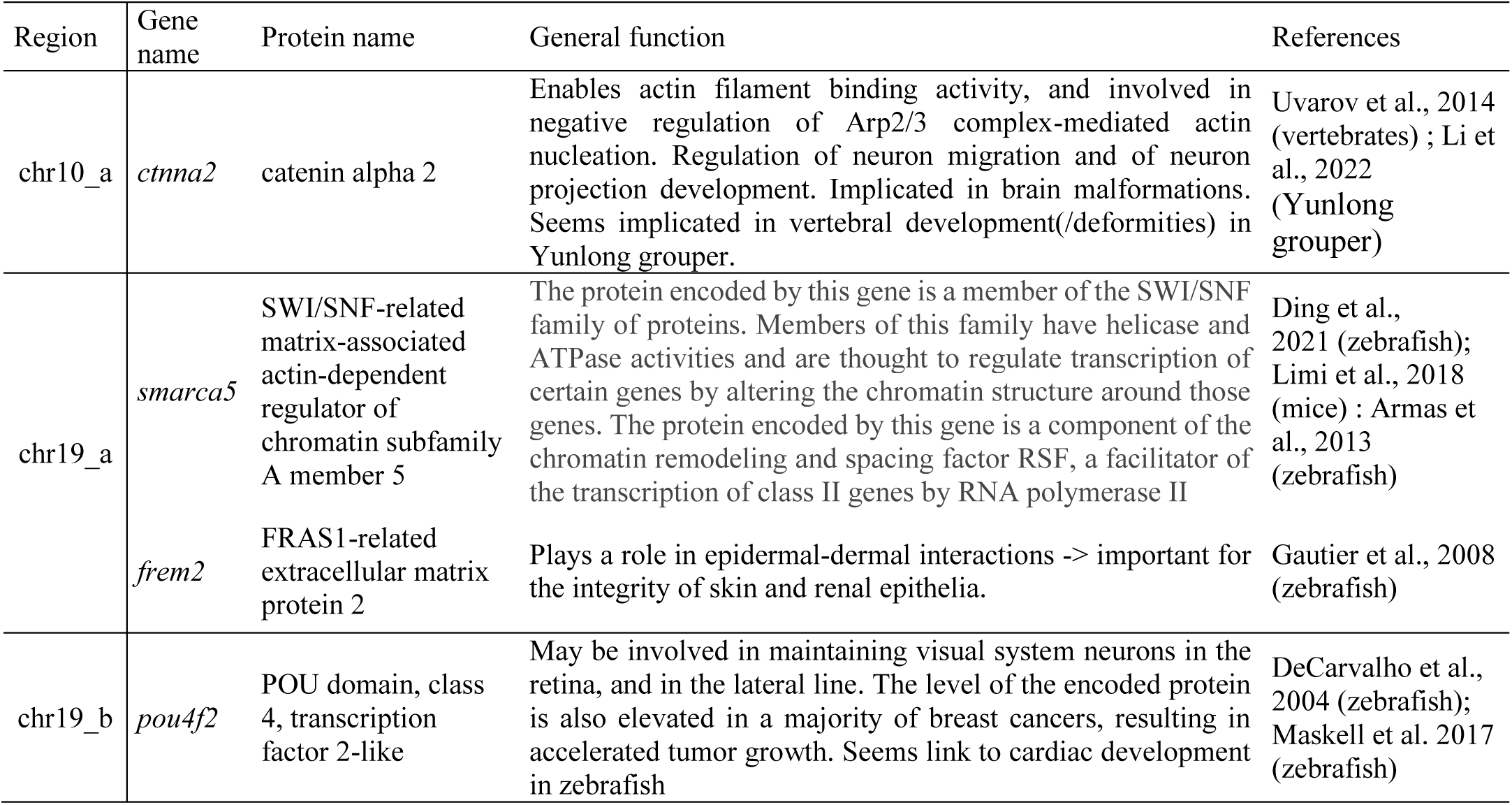
List and functions of the 4 genes annotated in three heterozygous regions (chr10_a, chr19_a, and chr19_b) shared by the four rainbow trout populations.

## 4 Discussion

The objective of our study was to detect signatures of selection into domestic rainbow trout. To reach that goal we studied four genetically distinct populations coming from different locations either in France or in the North-West of the USA. We used two different approaches, ROH and EHH, to detect the genomic regions shared by all populations using a HD SNP. We were able to detect 9 very conserved regions and 4 hotspots of polymorphism, corresponding to 253 and 29 annotated genes, respectively.

### 4.1 Genetic structure

First, we described the genetic structure of the populations under scrutiny. The three French lines were moderately differentiated with Fst ranging from 0.10 to 0.12. These estimations were congruent with those computed by D’Ambrosio et al.’study (2019) with the same populations that ranged between 0.09 and 0.14, but were estimated using a 38K SNPs array. These moderate differences between the 3 French populations were consistent with the PCA we performed and the history of these populations with a partly common INRAE origin (D’Ambrosio et al., 2019). This trend is shared between European populations with for instance an average Fst of 0.13 between 12 European rainbow trout strains (Gross et al., 2007) or 0.12 among 9 Norwegian populations (Glover, 2009). Similarly, US farmed populations are also weakly to moderately differenciated with an average Fst of about 0.09 (Silverstein et al., 2004) or 0.13 (Liu et al., 2017) and pairwise values ranging from 0.06 and 0.16. We observed a similar pattern in the present study with the HA population that consisted in samples from 5 locations, which all clustered together in the PCA. Reversely, we observed a large differentiation between our French and US populations revealed by large Fst values (0.27-0.29). This is likely the result of numerous factors, including selection, genetic drift and absence of gene flow between these very geographically distant populations. In addition, the European farmed populations originated from Californian domesticated strains, that have been shown to differ from strains of North-Western USA (Stanković et al., 2015). We found 34 haplotypes distributed over 21 chromosomes that differed between the American pooled population (HA) and all French populations (Supplementary information S1).

Due to the moderate to large differentiation between the 4 populations, the conserved regions across all populations are likely to be the result of ancient natural selection traces.

### 4.2 Comparison of methods to detect common signatures of positive selection

We used a double check of positive selection traces in the genome by using both ROH and EHH approaches. However, for each population, only a few regions were identified by both methods. These regions detected by more than one method represent stronger evidence of selection signatures since outlier markers detected by various genome scan methods help to uncover true selection signatures by reducing the number of false positives.

Even if both methods evaluate the homozygous large stretches in the genome, iHS also considers information based on haplotypic version and linkage disequilibrium from a core SNP. ROH approach detects homozygous regions regardless of their haplotypic versions, contrary to iHS. Thus, it may detect a signature of positive selection even if various haplotypes were present at the homozygous state in the population. In addition, while the ROH approach only detects the homozygous large stretches (at least 500 kb in the present study), iHS can detect small regions under positive selection as the only limitation in EHH region size is based on a threshold value for a minimum LD (0.10). Consequently, the sizes of the detected homozygous region varied between 1,065 kb and 2,857 kb based on ROH metrics and between 1,000 kb and 1,600 kb with iHS statistics.

The high number of regions (55, 69, 73, and 362) detected by iHS in our study was consistent with numbers detected in either Atlantic salmon (López et al., 2019) or cattle (Saravanan et al., 2021). However, these two previous studies used a lower threshold than ours (log(p-value) = 3 and 2, respectively vs 4 in the present study). Lower numbers of regions were previously detected by iHS in rainbow trout by Cádiz et al. (2021) and in Coho salmon by López et al. (2021). We speculate that these differences in the numbers of detected signals may be linked to the lower density of SNPs they used in both studies (57K or 200K chip versus 665K chip for our study) and the subsequent lower ability to detect LD and haplotypes at fine scale. Indeed, in the Chilean rainbow trout study (Cádiz et al., 2021), only one signal of positive selection was detected by iHS located at 6.398-14.936 Mb on chromosome 20 of the Swanson reference genome, which corresponds to the region 7.488-16.111 Mb on chromosome 20 of the Arlee reference genome. Nevertheless, we also detected by iHS signals of selection in each of our four studied populations, located at 10.5-16.5 Mb for LB, 11.2 – 13.3 Mb for LC, 13.0 – 14.2 Mb for SY and 12.3-13.2 Mb for HA (Supplementary Informations S6 to S9 for LB, LC, SY, and HA, respectively). Thus, all these signals were consistent with the larger region identified by Cádiz et al. (2021).

A common putative selection signature located at 13.0-13.2Mb could also be shared by all studied populations. In this 200kb-region, we observed at least one iHS value over |2.5| for LB, LC and SY lines, but not for HA population. In this region of 200 kb on chromosome 20, six genes were identified (*lgi1, noc3l, plce1, slc35g1, fra10ac1, tbc1d12*). Among these genes, Cádiz et al. (2021) identified two candidates genes associated with domestication, *noc3l* (nucleolar complex protein 3 homolog) and *plce1* (1-phosphatidylinositol 4,5-bisphosphate phosphodiesterase epsilon-1). Both are related to early development traits in zebrafish (*noc3l*: Walter et al., 2009; *plce1*: Zhou et al., 2009).

### 4.3 Biological functions of genes under positive or balancing selection

Among the 282 genes in the 13 regions detected under either positive or balancing selection, most genes seem to play essential roles in fitness as expected with such a dataset comprising both European and US populations. They are related to all main biological functions (genome stability, cell organization, neuronal and embryonic development, energy metabolism, growth, reproduction, and immunity). All identified biological functions were already described in other studies of signatures of selection in farmed rainbow trout (Cádiz et al., 2021) and other domesticated species (López et al., 2018, 2018; Naval-Sanchez et al., 2020; Baesjou & Wellenreuther, 2021; Signer-Hasler et al., 2022).

#### 4.3.1 Hotspots of heterozygosity and balancing selection for fitness traits

In livestock species, many variants under balancing selection are known to improve performance in heterozygote state but cause defect in homozygous state (Hedrick, 2015; Georges et al., 2019). However, in such cases of balancing selection, there is generally only one homozygous state, which is deleterious at a locus level, while the alternative homozygous state is observed in the population. In our study, we highlight four regions potentially involved in balancing selection for which we observed a lack of any kind of long streches of homozygosity. Even if these regions are extremely heterozygous, the proteins associated with these genes are highly conserved among vertebrates (Supplementary information S17). Many processes may explain these surprising observations at first glance. First of all, these regions may concentrate polymorphism in non-coding parts of the genome. Polymorphism in intronic regions may promote various proteins by allowing alternative splicing. We may also observe an excess of synonymous polymorphism in exons without effects on proteins. Further analyses must be conducted to better understand the mechanisms underlying the maintenance of extreme polymorphism, whether to validate the hypothesis of balancing selection or the existence of high mutation and recombination rates in these specific regions.

In the heterozygous region chr10_a, the gene *ctnna2 (*Table 8) enables actin filament binding activity and is involved in the regulation of neuron migration and neuron projection development. Thus, *ctnna2* plays an essential role in brain development among vertebrates (Uvarov et al., 2014). In yonlong grouper, *ctnna2* seems implicated in vertebral development, because significantly differentially expressed between normal and fish with lordosis (Li et al., 2022). In mice, a homozygous for a mutation of *ctnna2* reduced body weight, male fertility, and induced brain abnormalities (hypoplastic cerebellum, abnormal foliation pattern, ectopic Purkinje cells, and abnormal pyramidal cells in the hippocampus). While the protein associated to this gene is highly conserved among vertebrates (Uvarov et al., 2014; Supplementary information S17), the gene exhibits a strong polymorphism in all the four studied rainbow trout populations. However, a large part of its polymorphism is located in one intronic region (intron 6-7) of *ctnna2*. In the zfin database, five transcripts of this gene were identified (three mRNA and two non-coding RNA). We hypothezise that the polymorphism in the intronic region of *ctnna2* is essential for alternative splicing.

In the heterozygous region chr13_a (Supplementary information S16), *mmd* and *map2k4* are identified as highly conserved across vertebrates (Supplementary information S17). The gene *mmd* plays an important role in maturing macrophages, which is essential for immune response as observed in mice (Lin et al., 2021). The gene *map2k4* is implicated in a variety of cellular processes (proliferation, differentiation, transcription regulation, development), seems to play a role in liver organogenesis and embryonic development during gastrulation, as demonstrated by morpholino-mediated knockdown in zebrafish (Seo et al., 2010), and implicated in immune response in yellow catfish (Zheng et al., 2022). The inflammatory process in immune response seems linked to the polymorphism of the *map2k4* gene, which is consistent with our hypothesis of balancing selection, and more precisely potential ancestral trans-species polymorphism in this genomic region (Gu et al., 2016; Fijarczyk & Babik, 2015). Trans-species polymorphism is a crucial evolutionary mechanism for sharing adaptative genetic variation across taxa (Klein et al., 1998). The study of this mechanism has primarly concentrated on major histocompatibility complex genes, but a few studies described this process for other immune genes (Ferrer-Admetlla., et al., 2008; Leffler et al., 2013; Těšický & Vinkler, 2015). Maintaining genetic diversity in regions related to the immune system may be essential to resilience against various pathogens. In addition, this region of chromosome 13 has been recently detected as a significant QTL playing a role on resistance to temperature (Lagarde et al., 2022).

In the heterozygous region chr19_a (Table 8), the protein encoded by *smarca5* is a component of chromatin remodeling and spacing factor RSF, a facilitator of the transcription of class II genes by RNA polymerase II (zebrafish: Armas et al., 2013; Ding et al., 2021; mice: Limi et al., 2018). The protein is highly conserved among vertebrates (Supplementary information S17), which is consistent with its essential role thought to regulate the transcription of many genes by altering the chromatin structure around those genes. In the same region chr19_a, *frem2* codes for an extracellular matrix protein required for maintenance of the integrity of skin and renal epithelia in zebrafish (Gautier et al., 2008). This protein is moderately conserved across vertebrates (Supplementary information S17). In a study searching for genomic regions with ancestral trans-species polymorphism shared between humans and chimpanzees (Leffler et al., 2013), *frem*3, an important paralog of *frem2,* was identified under balancing selection. However, further studies should test the hypothesis of trans-species conservation of *map2k4* and *frem2* genes that may help to understand the various cellular processes in which the gene is implicated.

In the heterozygous region chr19_b (Table 8), *pou4f2* protein is highly conserved among vertebrates (Supplementary information S17) and is a tissue-specific DNA-binding transcription factor involved in the development and differentiation of specific cells. It maintains the visual system neurons in the retina and the lateral line (DeCarvalho et al., 2004) and seems also related to cardiac development in zebrafish (Maskell et al., 2017).

#### 4.3.2 Hotspots of homozygosity and positive selection for essential biological functions

##### 4.3.2.1 Regions and genes involved in cellular and nuclear organization, and embryonic development

In homozygous region chr2_a, three genes plays important roles in embryonic development and then fitness (*cep162*, *tsnare1*, *mrap2*, Table 6). In the homozygous region chr2_b (Table 6), the gene *pbx1* (pre-B-cell leukemia transcription factor 1) is related to early development in zebrafish (Teoh et al., 2010). Mutations in this gene generally cause major malformations, which seem to play an essential role in survival in various species (zebrafish: Teoh et al., 2010; mouse: Selleri et al., 2001; human: Le Tanno et al., 2017). It was detected under positive selection in a Chilean farmed rainbow trout population (Cádiz et al., 2021). However only moderate percent identity (> 65%) is observed between *pbx1* proteins across vertebrates.

In the homozygous region chr15_a, many genes *(chn1, atp5mc1, zc3h15, nid2 and brca2)* were playing essential roles in cell functioning and early development (Table 6). However only two of them were highly conserved among vertebrates (> 85%; *chn1* and *atp5mc1*). The gene *atp5mc1* is a crucial gene for mitochondrial cristae morphology, and plays important roles in metabolic processes associated to growth (Table 6; Palmer et al., 2011; Miller et al., 2019; Wang et al., 2020). In zebrafish, a morpholino knockdown of *chn1* reveals its crucial role in early development, revealing severe abnormalities (development of round somites, lack of yolf extension, and kinkled posterior notochord) (Leskow et al., 2006).

Three genes located in close vicinity in region chr16_a (between 46.42 and 46.53 Mb; Supplementary information S11), *samd10* (sterile alpha motif domain-containing protein 10-like), d*najc5* (dnaJ homolog subfamily C member 5-like), and *tpd54* (tumor protein D54) were also detected in close chromosomal vicinity and under positive selection in ten modern goat breeds and one wild Bezoar goat (Signer-Hasler et al., 2022). This cluster of genes has a significant role in survival and cellular processes (Table 6). In addition, in this region chr16_a, the protein of the gene *magi2* (membrane-associated guanylate kinase, WW and PDZ domain-containing protein 2, Table 6) plays a vital role in embryogenesis in zebrafish (Borah et al., 2016). The gene *magi2* was also identified under positive selection in a domesticated sheep breed compared to the wild Asiatic mouflon (Cumer et al., 2021).

##### 4.3.2.2 Regions and genes involved in neural and brain development, and behaviour

In total, we identified 7 genes as primarly associated to neural and brain development in both regions detected under positive selection (*tsnare1, cdk14, brsk2a, auts2, brd2, znf135,* and *grxcr1*). Some genes (*brsk2a, znf135, grxcr1, auts2; Table 6*), related to brain development may induce behavior modifications in farmed animals, that may be related to domestication processes (Pasquet, 2018; Milla et al., 2021; Deng et al., 2022; Liu et al., 2022). This is in line with Żarski et al. (2020) study showing that domestication modulates gene expression involved in neurogenesis.

In particular, the gene *auts2* gene was previously identified under positive selection both in cattle (Consortium, bovine Hapmap, 2009) and in domesticated Atlantic salmon populations from Canada and Scotland compared to their wild Atlantic salmon counterpart (López et al., 2018). The gene *znf135* was also detected under positive selection in a farmed population of Atlantic salmon compared to a wild-type population (Gutierrez et al., 2016). The gene *grxcr1* was detected under positive selection in the Tharparkar cattle (Saravanan et al., 2021). It strongly suggests that all these genes play a key role in domestication processes and may act on essential behaviours in both terrestrial and aquatic farmed animals.

##### 4.3.2.3 Regions and genes involved in growth metabolism

Genes related to growth metabolism were only identified in four regions under positive selection, none of them were detected in high heterozygosity regions.

In the homozygous region chr2_a (Table 6), the protein *mrap2* (melanocortin-2 receptor accessory protein 2A) is associated to growth. A lack of this gene exhibit severe obesity in many species (human, zebrafish, rodent: Liu et al., 2013; sea lamprey: Zhu et al., 2019; snakehead: Wang et al., 2021). Yoshida et al. (2017) detected a growth-QTL in Atlantic salmon and considered *mrap2* as a candidate gene for growth up to 25 months. In addition, *mrap2* was identified in the Chilean farmed rainbow trout population as under positive selection (Cádiz et al., 2021). A QTL related to sea lice resistance in rainbow trout (Cáceres et al., 2021) was also detected in the region chr2_a (located from 10.43 Mb to 11.81 Mb of the Swanson reference genome, which corresponds to 26.69 Mb – 28.07 Mb of the Arlee reference genome). Cáceres et al. (2021) explained that having a high potential for growth seem essential for sea lice resistance. Indeed, proteomic investigations allow to establish a link between growth and immune function in salmonids (Causey, 2018).

In homozygous region chr2_b (Table 6), the *col9a2* (collagen alpha-2(IX) chain) gene is a component of cartilage and seems also related to growth (Xu et al., 2022). This gene is detected under positive selection in a Scottish farmed population of Atlantic salmon (López et al., 2018). In addition, the gene *scap* (sterol regulatory element-binding protein cleavage-activating protein) was already identified under positive selection in six farmed Pacific white Shrimp populations (Wang et al., 2022).

In the homozygous region chr2_d (Table 6), the gene *igf-1α* (insuline like growth factor receptor 1a) plays an important role in growth and transformation events. In salmonids, expressions of *igf-1α* and growth hormone were demonstrated to be modified between domesticated and wild populations of rainbow trout and coho salmon (Tymchuk et al., 2009). The same observation was made with a higher expression of *igf-1α* between larvae from domesticated spawners than larvae from wild spawners of the Eurasian perch (Palińska-Żarska et al., 2021). In addition, *igf-1* was also observed as a marker of domestication in dogs (Wayne & vonHoldt, 2012).

In the homozygous region chr16_a, the *emilin-3a* gene (elastin microfibril interfacer 3a, Table 6) plays a role in extracellular matrix organization and elastic fiber formation. Its gene expression is related to embryonic development and involved in muscle fiber development in zebrafish (Milanetto et al., 2008). *Emilin-3a* had already been identified as under positive selection in one population of F2 Australian snapper farmed population compared to the first generation (F1) of domestication of a wild population (Baesjou & Wellenreuther, 2021). Thus, this signature of selection can be considered as a result of the domestication process.

All identified growth-related genes seem associated with domestication. This assertion is confirmed for five genes (*mrap2*, *col9a2*, *scap*, *igf-1α, emilin-3*) that were also identified under positive selection in various farmed populations with favorable alleles linked to better growth phenotypes (Table 6).

##### 4.3.2.4 Regions and genes involved in reproduction

Very few genes directly related to reproduction traits were only identified in highly homozygous regions.

In the homozygous region chr2_b, the *brd2* (bromodomain-containing protein 2, Table 6) gene is implicated in several biological process (see section 4.4.1.). It seems related to oogenesis and egg-to-embryo transition in zebrafish (DiBenedetto et al., 2008), which is consistent with a QTL detected for egg size in this region in rainbow trout (D’Ambrosio et al., 2020). Moreover, it seems that *brd2* is involved in spermatogenesis or folliculogenesis, as demonstrated in situ on mice cells (Rhee et al., 1998). Khendek et al. (2017) compared the reproductive performances (egg size, gonadal histology, hormonal levels) between domesticated and F1 with wild broodstock of Eurasian perch populations. They showed that domestication may have increased the oocyte diameter and the level of 17β-Estradiol, but decreased the embryo survival of domesticated fish. This gene was also identified under positive selection in a selected Canadian population of Atlantic salmon (López et al., 2018).

In the homozygous region chr15_a (Table 6), the gene *zp4* has already been identified under positive selection in a farmed Scottish population of Atlantic salmon compared to a wild population (López et al., 2018), and may be related to domestication process.

##### 4.3.2.5 Regions and genes involved in immunity

Magris et al. (2022) observed that regions under positive selection revealed an enrichment of KEGG terms related to viral infection in farmed brown trout. However, it should be noticed that in our study, few genes related to immune function were identified and no enrichment in immune terms was observed in GO analysis.

Genes related to immune function were mainly identified in three different regions detected as putatively under positive selection for two of them and under balancing selection for the last one.

In the homozygous region chr2_b (Table 6), genes *tnfaip8l2b* (tumor necrosis factor, alpha-induced protein 8-like protein 2 B) and *atg5* (autophagy protein 5) are related to immune functions. Note that atg5 is well conserved across vertebrates (> 80%).

In the region chr15_a, the gene *zc3h15* (Table 6) seems to have an inhibitory effect on HIV-1 replication and then on HIV infection in vitro (mice cells) (Capalbo et al., 2010).

A last gene in the region chr16_a, the gene *atp1b3* (sodium/potassium-transporting ATPase subunit beta-1-interacting protein 3) was also identified under positive selection in farmed Atlantic Salmon (Naval-Sanchez et al., 2020). In the Senegalese sole, atp1b3a and atp1b3b paralogs have been hypothesized to be involved in response to low salinity (Armesto et al., 2015). In addition, this gene is involved in some immune responses. It was shown in cell culture study, that *atp1b3* inhibits hepatitis B virus replication via inducing NF-kappa B activation (human: Zhang et al., 2021) and is involved in numerous viral propagation such as HIV and EV71 (Zheng et al., 2020) in cell culture experiments.

### 4.4. Conclusion

To sum up, we identified 13 regions under selection with numerous genes strongly involved in essential biological functions. By identifying signatures of selection shared by our four studied populations, we have focused our detection on regions related to ancient evolutionary processes that are essentially important for species survival. We only identified nine homozygous regions presumably under positive selection and four heterozygous regions putatively under balancing selection in four different rainbow populations. While common homozygous regions may be associated with important biological functions underlying both fish fitness and domestication, the heterozygous regions seem mainly linked to fitness functions (cell organization, embryonic development, and immunity) which are involved at different developmental stages or to cope with various pathogens or abiotic stressors. Maintaining genetic diversity in these regions could be essential for the species survival.

This study allows us to confirm the importance of a large set of 17 genes already detected as under positive selection in previous studies, among which 10 genes were identified in fishes (*auts2, atp1b3, zp4, znf135, igf-1α brd2, col9a2, mrap2, pbx1* and *emilin-3*). We also identify new promising candidate genes as important for rainbow trout fitness. In our opinion, this study substantially increases knowledge of evolutionary processes and helps to determine the genomic location and the nature of the genetic variation that must be maintained in rainbow trout populations for domestication and selection purposes.

## Acknowledgements

We thank the two breeding compagnies “Viviers de Sarrance” and “Milin Nevez” that allow us to use their HD genotypes to perform the study. This study was partly supported by the European Maritime and Fisheries Fund and FranceAgrimer (Hypotemp project, n◦ P FEA470019FA1000016).

## Data Accessibility and Benefit-Sharing

Restrictions applied to the availability of the data that support the findings of this study, which were used under license and so are not publicly available. The data can be made available for reproduction of the results from Florence Phocas on request via a material transfer agreement and with permission of the two breeding companies “Viviers de Sarrance” (Sarrance, France) and “Milin Nevez” (Plouigneau, France).

## Author contributions

Katy Paul: Investigations, Methodology, Formal analysis, Writing - Original Draft; Gwendal Restoux: Conceptualization, Methodology, Draft Reviewing; Florence Phocas: Supervision, Conceptualization, Methodology, Investigation, Formal analysis, Resources, Writing - Original Draft.

## Appendices

**Supplementary figure 1.**
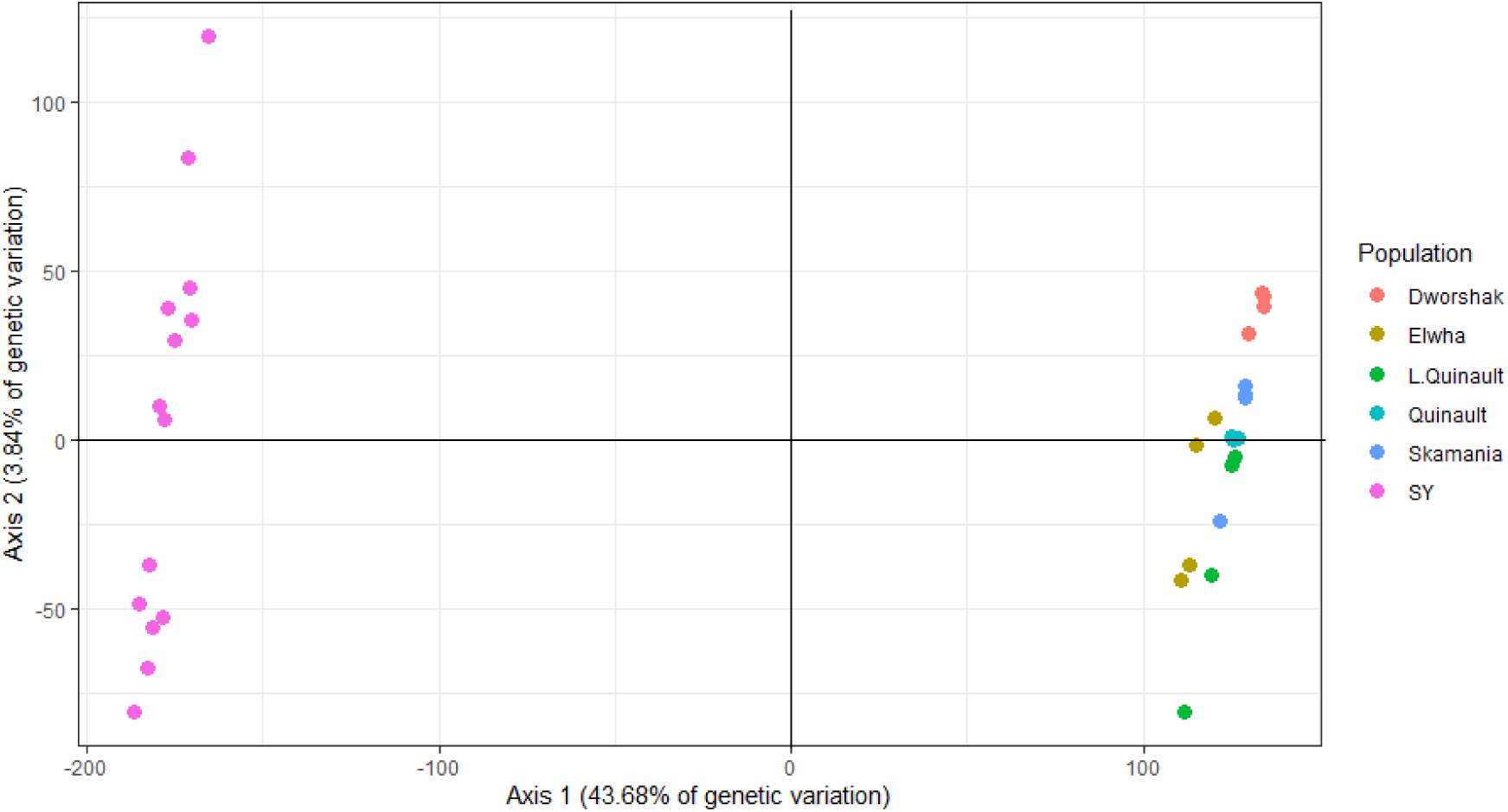
Principal component analysis (PCA) of the genetic diversity of SY, and HA sub-populations based on 546,903 SNPs.

<u>File:</u> *Supplementary_Tables.xlsx*

